# Osteocalcin binds to a GPRC6A Venus fly trap allosteric site to positively modulate GPRC6A signaling

**DOI:** 10.1101/2022.02.15.480526

**Authors:** Rupesh Agarwal, Min Pi, Ruisong Ye, Micholas Dean Smith, Jeremy C. Smith, L. Darryl Quarles

## Abstract

GPRC6A is a member of the Family C G-protein coupled receptors that is activated by cations, L-amino acids, the osteocalcin (Ocn) peptide, and testosterone. GPRC6A functions as a master regulator of energy metabolism and sex hormone production. Based on homology to the related receptors mGluR5 and CaSR, GPRC6A’s multiple ligand specificity is likely based on an orthosteric ligand binding site in the bilobed Venus fly trap (VFT) domain together with two positive allosteric modulator (PAM) sites, one in the VFT and the other in the 7TM domain. Here, we show that Ocn acts as a PAM for GPRC6A by binding to a site in the VFT that is distinct from the orthosteric site for calcium and L-amino acids. In agreement with this finding, alternatively spliced GPRC6A isoforms 2 and 3, which lack regions of the VFT, and mutations in the predicted Ocn binding site, K352E and H355P, prevent Ocn activation of GPRC6A. These observations provide a structural framework for understanding the ability of multiple distinct classes of compounds to activate GPRC6A and set the stage to develop novel small molecules to activate and inhibit this receptor.

## Introduction

Osteocalcin (Ocn) is a 49 residue protein in humans that is mainly produced by osteoblasts in bone. A vitamin K-dependent carboxylated form of Ocn comprises much of the non-collagenous bone protein matrix, while an undercarboxylated form (unOcn) is released into the circulation to function as a hormone. unOcn regulates energy metabolism through direct effects on metabolic functions in multiple target tissues and by indirect effects mediated by an ensemble of metabolically active hormones whose release is stimulated by unOcn (1–5). Indeed, unOcn is reported to stimulate insulin secretion and β-cell proliferation in the pancreas (5), fibroblast growth factor 21 (FGF-21) release and glucose and fat metabolism in liver hepatocytes (6), interleukin 6 (IL-6) secretion and glucose utilization in skeletal muscle myocytes (7), adiponectin release and lipolytic activity in adipocytes in white fat (8), testosterone production from testicular Leydig (2,6), and glucagon-like peptide 1 (GLP-1) secretion from gastrointestinal enterocytes (9,10).

The peripheral metabolic effects of Ocn are mediated through activation of GPRC6A, a member of the Family C G-protein coupled receptors (5,11). GPRC6A is activated by several orthosteric ligands, including basic L-amino acids (such as L-Lys, L-Arg, and L-ornithine) and divalent cations (such as Ca^2+^ and Mg^2+^). Also, positive allosteric modulators (PAMs) of GPRC6A include testosterone, NPS-568, and di-phenyl and tri-phenyl compounds (12,13). Gallate and epigallocatechin 3-gallate (ECGC), which are natural products in green tea, respectively activate and inhibit GPRC6A (14). The structural basis for GPRC6A interaction with multiple ligands is incompletely understood.

Family C GPCRs are characterized by their similarity in 7 transmembrane (7TM) domain sequences, a necessity for dimerization to function, and in most cases, the existence of a large N-terminal ligand-binding Venus flytrap (VFT) domain (15). Recently, high resolution structural data of the related mGluR5 (16) and CaSR (17–20), have identified how various ligands and drug molecules target the VFT and 7TM. For example, mGluR5 has the orthosteric binding site for L-glutamate in the VFT, and two positive modulating allosteric (PAM) sites, one located in the VFT and the other in the 7TM (16). The allosteric site located in the VFT was discovered by a nanobody, Nb43, that binds to Helix L and the L-M loop of mGluR5 to potentiate orthosteric agonist binding and receptor activation. Similarly, elucidation of the 3D structure of CaSR identified in the VFT an orthosteric Ca^2+^ ion binding site that acts as a composite agonist with L-amino acids, such as L-tryptophan, to stabilize the closure of the active VFT (17). CaSR also has an allosteric site at the dimeric interface of the VFT (proximal to residue D248) to which the peptide etecalcitide binds and acts as a positive allosteric modulator (PAM). In CaSR, additional small molecule drugs act as PAMs (e.g., calcimimetics), or negative allosteric modulators (NAMs) (e.g., calcilytic) via a site in the 7TM (17–20).

Based on this new structural information, we developed a structural model for the activation of GPRC6A by orthosteric and allosteric ligands. We investigated Ocn activation of GPRC6A isoforms with deleted segments of the extracellular domain, mapped the peptide fragments of Ocn necessary for activation of GPRC6A, and examined if Ocn peptides bind to the VFT and modulate GPRC6A function.

## Results

### Structural modeling of GPRC6A isoforms and L-Arginine orthosteric ligand binding

GPRC6A exists as three isoforms: isoform 1 represents the full-length receptor, and two alternatively spliced receptor isoforms 2 and 3 are characterized by respective deletions of exon 3 and exon 4 segments that encode regions of the VFT domain (21) (**Figure 1A**). To identify regions that differ between the isoforms, we aligned the protein sequences of isoforms 1, 2 and 3 of GPRC6A (**Figure 1B, S1**). The TM domains of these three isoforms have 100% sequence identity, but isoforms 2 and 3 are missing residues 271-445 and 446-516 in the extracellular domain, respectively.

**Figure 1.**
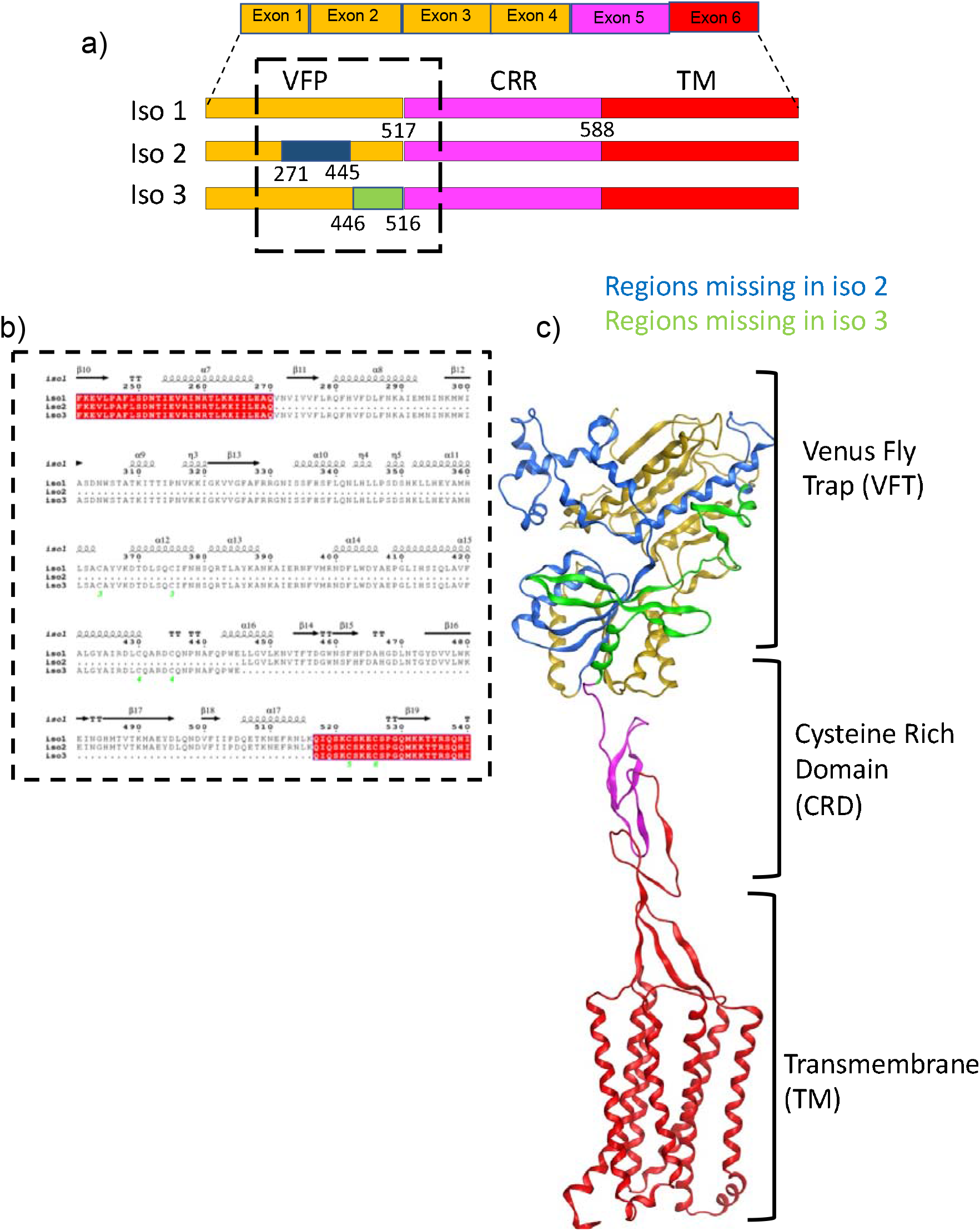
Sequence and structure of GPRC6a. a) Placement of exons and its location in GPRC6A isoform 1, 2 and 3 structure; b) Sequence alignment of Venus flytrap (VFT) domain showing missing regions of isoform 2 and 3; Structure model of GPRC6A showing the three domains and the missing region of isoform 2 and 3.

To derive the structure of GPRC6A, we used AlphaFold2 (22–25) to model the full-length isoform 1 (**Figure 1C**). AlphaFold2 is an AI-based technique that derives close-to-experimental accuracy in protein structure prediction. These calculations confirmed that GPRC6A has 3 domains: TM, CRD and VFT, similar to the related mGluR5 and CaSR Cryo-EM structures. The residues deleted in the isoforms 2 and 3 correspond to different segments of the VFT domain. In the case of isoform 2, a large segment of the VFT domain structure (~ 174 residues) is deleted, whereas a relatively small region (~ 70 residues) is missing in isoform 3 (**Figure S1**).

In isoform 1, we identified an orthosteric ligand site near residues Tyr 148, Ser 149, Thr 172, Asp 303 in the VFT based on structure superimposition with mGluR5 and CaSR (**Figure S2a and S2b**). In contrast, in isoform 2, the orthosteric ligand binding site predicted based on aligning orthosteric bound structures of mGluR5 and CaSR structures is missing, foretelling a loss-of-function (*vide infra*) (**Figure S2c**). The predicted orthosteric ligand binding site is retained in isoform 3, and hence this isoform is predicted to be responsive to orthosteric ligands (*vide infra*). To test these predictions, GPRC6A isoforms 1, 2, or 3 were transfected into HEK-293 cells and receptor activation was measured by cAMP accumulation in response to the orthosteric ligand L-Arg in the presence of 1 mM calcium. Isoform 2 resulted in loss of L-Arg stimulation of cAMP, whereas isoform 3 retained responsiveness to L-Arg activation, similar to isoform 1 (**Figure 2A**).

**Figure 2.**
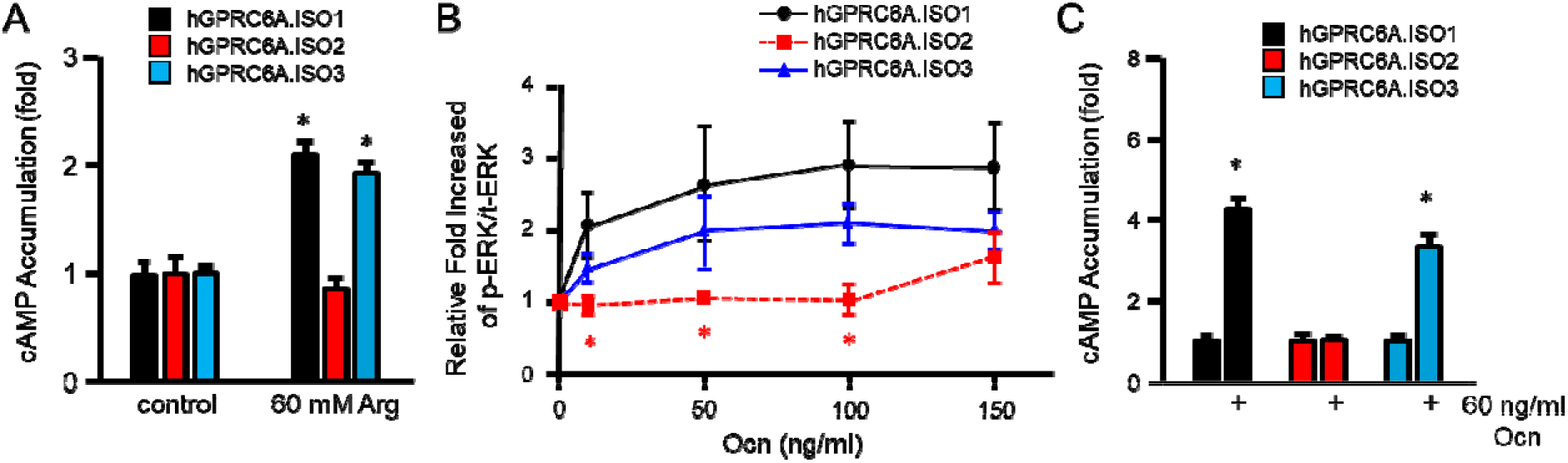
Activities of GPRC6A isoforms on response to Ocn. A. Comparison of effects of L-Arg on GPRC6A isoforms mediated cAMP accumulation. Comparison of dose-dependent effects of Ocn on GPRC6A isoforms mediated ERK activation (B) and cAMP accumulation (C). HEK-293 cells were transfected with cDNA plasmids of GPRC6A isoform 1, 2 or 3 for 48 hours, after incubated in Dulbecco’s modified Eagle’s medium /F-12 containing 0.1% bovine serum albumin quiescence media for 4 hours, then exposed to L-Arg or Ocn at indicated concentrations for 15 minutes for ERK activation, or 40 minutes for cAMP accumulation details as described under “Methods”. * indicates a significant difference from control and stimulation groups at p<0.05.

### GPRC6A isoforms 2 and 3 have attenuated responses to Ocn

The location of the Ocn binding site in GPRC6A is still not well understood. To explore the role of the VFT in mediating the response to Ocn, we compared the ability of Ocn to activate the three GPRC6A isoforms (**Figures 2B and C**). In the presence of 1 mM calcium Ocn activated GPRC6A isoform 1 in a dose dependent manner, consistent with previous reports (5). GPRC6A isoform 2 exhibited an attenuated ERK response with a complete loss of cAMP activation in response to Ocn treatment. In contrast, GPRC6A isoform 3 also showed attenuation of Ocn-induced ERK stimulation but the cAMP response to Ocn persisted (**Figures 2B and C**). Taken together, loss-of Ocn responses in isoforms 2 and 3 suggest that the VFT is necessary for Ocn activation of GPRC6A and that the different isoforms bias ligand-induced coupling to G-protein signaling.

### Identifying Ocn amino acids necessary for GPRC6A activation

Ocn in humans is a 49 amino acid peptide. Initially, we used functional assays to refine the protein-protein interface between Ocn and GPRC6A VFT. We designed Ocn fragments (**Figure 3A**) (11,26) and compared their ability to activate the full length GPRC6A isoform 1 (**Figures 3B and C**) transfected into HEK293 cells. Ocn peptide activation was measured by assessing both ERK (**Figure 3B**) and cAMP (**Figure 3C**) signaling. Compared to the full length Ocn (1–49), all peptide fragments activated signaling in cell transfected with GPRC6A to some degree. However, we found that the decapeptide fragment of Ocn 20-29 (revcelnpdc) located in the middle of the protein and the C-terminal hexapeptide Ocn 44-49 (rfygpv) exhibited significantly higher activity (**Figure 3B and C**).

**Figure 3.**
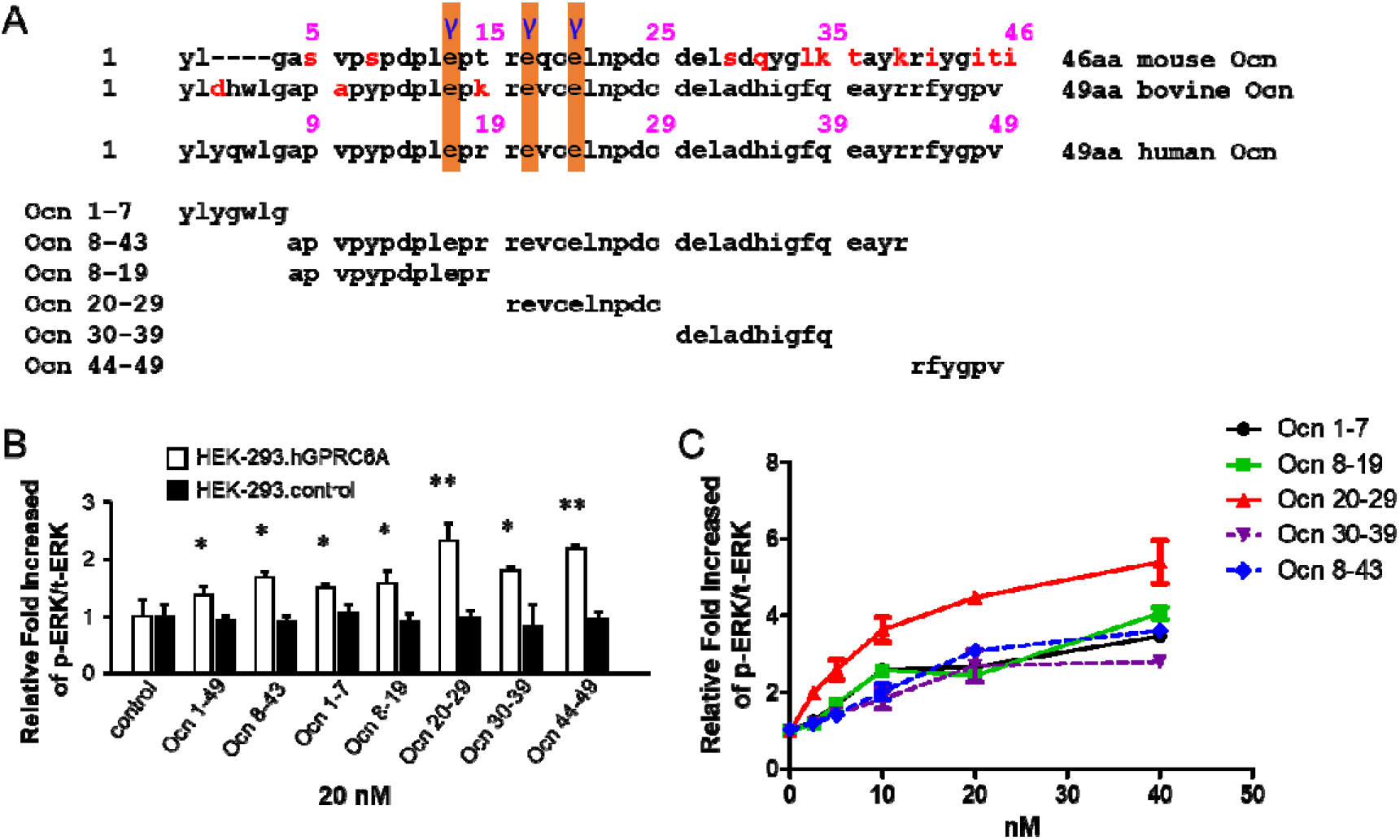
The activities of human Ocn peptide fragments. A, The sequence alignment of full length mouse, bovine and human Ocn (upper panel), and synthesized peptide fragments of human Ocn (bottom panel). Red alphabets indicate different amino acids compared to human Ocn sequence. “γ” indicates carboxylation sites in Ocn. B, Comparison of the activities of synthesized fragments of human Ocn (at 20 nM) by ERK phosphorylation in HEK-293 transfected with vector pcDNA3 (black bar) and HEK-293 cells transfected with pcDNA3-hGPRC6A cDNA (white bar). * and ** indicate a significant difference from control and stimulation groups at p<0.05 and p<0.01, respectively. C, Dose response of synthesized fragments of Ocn in HEK-293 cells transfected with pcDNA3-hGPRC6A cDNA. The ERK phosphorylation was measured 15 minutes for Ocn fragments at concentration as indicated at indicated stimulation in HEK-293 cells transfected with pcDNA3-hGPRC6A cDNA after 4 hours quiescence.

To further investigate the bioactivity of Ocn 20-29 fragment *in vivo*, we injected the synthetic Ocn 20-29 peptide (2 μM/kg) into wild type *Gprc6a*^+/+^ and *Gprc6a*^-/-^ mice by the intraperitoneal (IP) route (27), and collected blood and serum at 24 hours. We found that Ocn 20-29 at a dose of 2 μM/kg reduced blood glucose levels in wild-type mice by ~10% at 24 hours after intraperitoneal administration of 2 μM/kg, whereas the vehicle (saline) had no effect on blood glucose (Fig. 4A). Consistent with Ocn effects in stimulating IL-6 release from skeletal muscle (7), Ocn 20-29 at a dose of 2 μM/kg IP also resulted in a significant increase in serum IL-6 in wild type *Gprc6a*^+/+^ mice, but not in *Gprc6a* knockout (*Gprc6a*^-/-^) mice (Fig. 4C). Ocn 20-29 injection did not significantly stimulate serum insulin (Fig. 4B) or adiponectin (Fig. 4D) levels in wild-type mice at 24 hours at a dose of 2 μM/kg.

**Figure 4.**
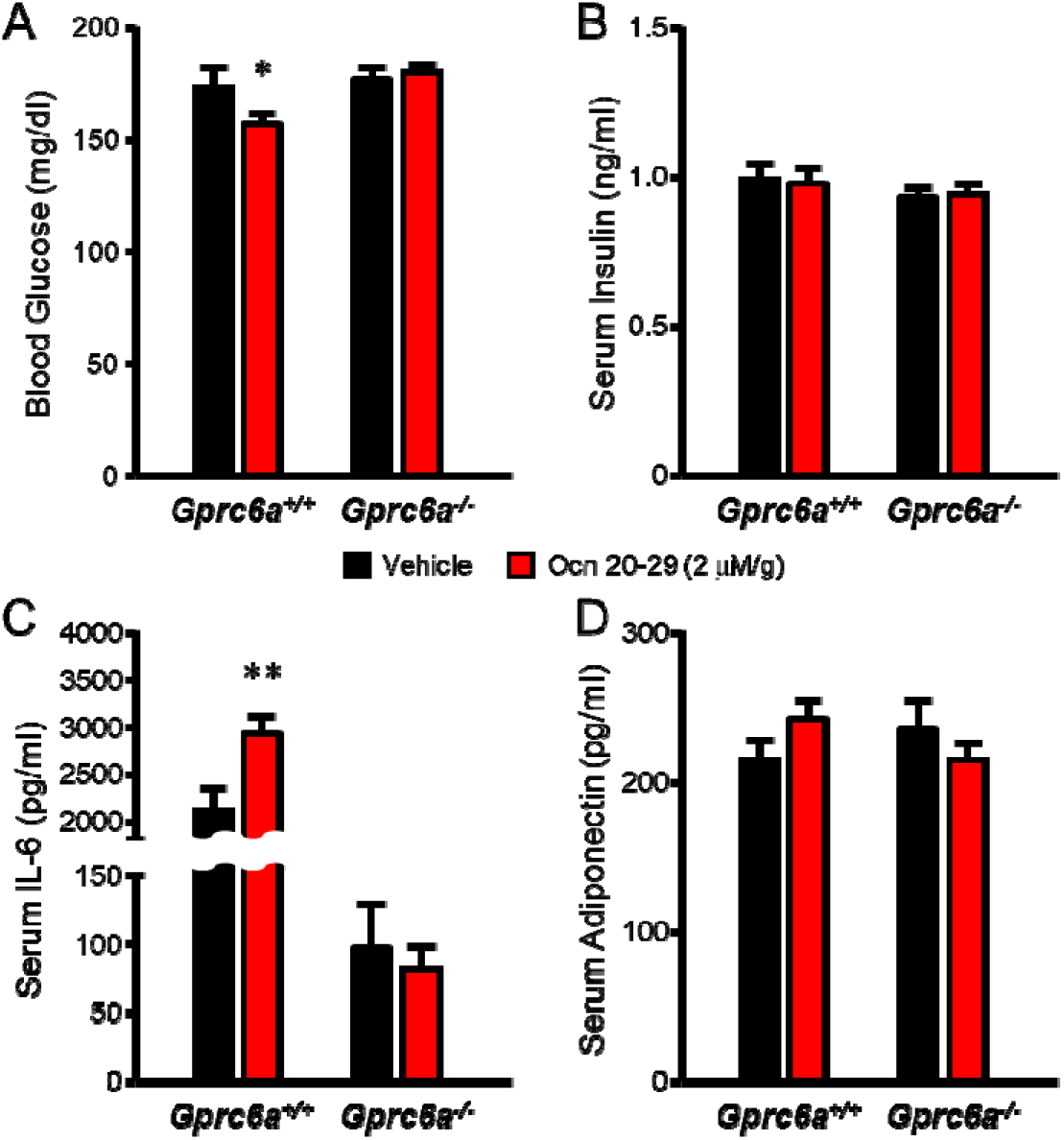
The effects of Ocn fragment, Ocn 20-29 on GPRC6A-mediated regulation of blood glucose and serum IL-6 levels in mouse. The biological activities of Ocn 20-29 were assessed 24 hours after intraperitoneal injection of 2 μg/kg the Ocn 20-29 peptide (red bar) or vehicle (saline, black bar) in 8 week-old *Gprc6a*^+/+^ or *Gprc6a*^-/-^ mice by measuring blood glucose (A), serum insulin (B), IL-6 (C), and adiponectin (D). Ocn 20-29 significantly decreased the blood glucose level, and increased serum IL-6 level compared to vehicle in wild-type mice. These response were lost in *Gprc6a*^-/-^ mice. We observed no differences in serum insulin or adiponectin levels in Ocn 20-29 and vehicle treated mice. * and ** indicate a significant difference from vehicle and Ocn 20-29 injection groups (n≥5) at p<0.05 and p<0.01, respectively.

### Protein-protein docking to generate GPRC6A-VFT: Ocn complex structure

As noted above, there are two PAM sites in the related receptors mGluR5 and CaSR, one in the 7TM domain targeted by small molecules and another in the VFT targeted by the protein nanobody (16) and the etecalcitide peptide (20). Specifically, the PAM function of Nb43 is mediated by a nanobody binding site near a helix in the VFT domain of the mGluR5 as defined by Cryo-EM (16). Aligning the structures of the allosteric ligand bound VFTs mGluR5 (PDB 6N4Y) and CaSR (PDB 7E6U, 7M3G) with our model of GPRC6A and its isoforms indicated that the dimeric interface found to bind etecalcitide and NAM in CaSR was invariant across the GPRC6A isoforms, while the binding pocket homologous to the mGluR5 nanobody PAM site was absent in GPRC6A’s isoform 2 (**Figure S3a**). We hypothesized that, given the loss of Ocn associated function in isoform 2, the pocket homologous to the mGluR5 binding site may serve as the binding site for Ocn in GPRC6A. Using this site as our guide, we used HADDOCK 2.4 (28) to generate a model of the GPRC6A-VFT: Ocn complex. The top 4 scoring structures within the calculated top cluster were then subjected to additional analysis and visual inspection.

Our analysis revealed that the GPRC6A-VFT forms multiple hydrogen bonds (a list is provided in **Table 1**) with Ocn in all the models. Residues 20-29 from the highest active Ocn fragment also form multiple hydrogen bonds with GPRC6A residues (Gln 19, Asp26, Gln381, Lys352) (**Figure 5**) and is also in proximity (within 4 Å) to other residues (Ala385, Arg382, Asp26, Asp349, Gln19, Gln22, Gln61, Glu356, Glu62, His351, His355, Pro20, Ser348, Ser350, Val28), which can form other potential electrostatic interactions. Moreover, the Ocn model shows that there is a disulfide bond between residues Cys 23 and Cys 29 (**Figure S3b**), both of which are present in the Ocn fragment. This is predicted to provide structural integrity to the fragment and would potentially fold similarly to the full length Ocn peptide. In analyzing the interactions between the GPRC6A-VFT and Ocn, we observe that two GPRC6A-VFT residues, Lys 352 and His 335, in the helix region (352–363) are common in all the models (**Figure 5, Table 1**). These residues form hydrogen bonds with Ocn residues Asp 28, Asp 30 and Glu 31 and are near several other residues present in 20-29 region of Ocn that was the most active fragment.

**Figure 5:**
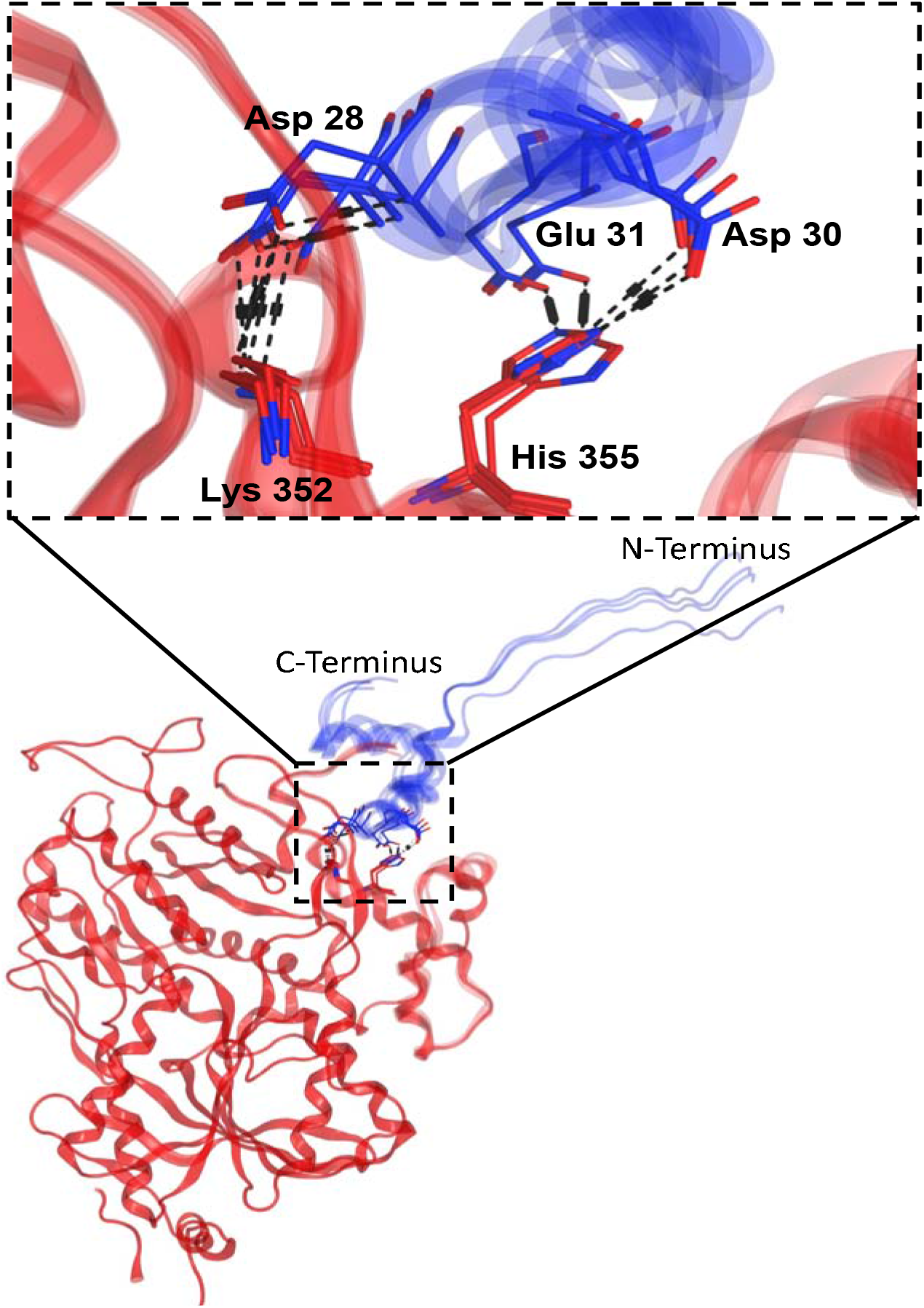
Superimposition of the top 4 models of GPRC6A’s VFT domain (in red) and Ocn (in blue) complex generated from HADDOCK. The key residue interactions present in the common in helix region (352–363) of GPRC6A’s VFT and residues 20-29 of Ocn.

**Table 1:** List of hydrogen bonds between GPRC6A-VFT and Ocn in best 4 structures from the top cluster from HADDOCK.

| GPRC6A residue id | Ocn residue id | Distance (Å) | Model number |
| --- | --- | --- | --- |
| Asn389 | Asp34 | 2.79 | 1 |
| Asp26 | Tyr46 | 2.73 | 1 |
| His355 | Asp30 | 2.93 | 1 |
| Lys352 | Asp28 | 3.31 | 1 |
| Ser348 | Glu31 | 2.67 | 1 |
| Asp26 | Asn26 | 2.81 | 2 |
| Asp26 | Phe45 | 3.83 | 2 |
| Asp26 | Tyr46 | 2.73 | 2 |
| Gln381 | Glu24 | 2.8 | 2 |
| Gln381 | Pro27 | 3.06 | 2 |
| His355 | Asp30 | 3.38 | 2 |
| His355 | Glu31 | 2.8 | 2 |
| Lys352 | Asp28 | 3.12 | 2 |
| Ser348 | Glu31 | 2.71 | 2 |
| Ala385 | Asp30 | 3.47 | 3 |
| Asn389 | Asp34 | 2.81 | 3 |
| Asn389 | His35 | 4.7 | 3 |
| Asp26 | Asn26 | 2.81 | 3 |
| Gln19 | Pro18 | 3.89 | 3 |
| Gln19 | Glu21 | 3.03 | 3 |
| Gln381 | Glu24 | 2.81 | 3 |
| His355 | Asp30 | 2.91 | 3 |
| Lys352 | Asp28 | 3.18 | 3 |
| Pro20 | Pro18 | 3.48 | 3 |
| Ser348 | Glu31 | 3.6 | 3 |
| Ala385 | Asp30 | 3.57 | 4 |
| Asp26 | Phe45 | 3.38 | 4 |
| Gln19 | Pro18 | 3.86 | 4 |
| Gln381 | Glu24 | 2.8 | 4 |
| Gln381 | Asp30 | 2.87 | 4 |
| His355 | Glu31 | 2.75 | 4 |
| Lys352 | Asp28 | 3.06 | 4 |
| Pro20 | Pro18 | 3.06 | 4 |
| Ser348 | Glu31 | 2.71 | 4 |

### Validation of the complex structure using single point mutagenesis

To test the above models, we mutated Lys352 or His355 in human GPRC6A, into Glu352 (mutant K352E) or Pro355 (mutant H355K), and also created double mutated Glu352/Pro355 (mutant K352E/H355P) GPRC6A. Mutants and wild-type hGPRC6A cDNAs were transiently transfected into HEK-293 cells and signaling was assessed in response to Ocn treatment.

We confirmed that mutants K352E, H355P, K352E/H355P, and WT mGRPC6A proteins were equally expressed, as assessed by Western blotting using a Myc antibody, which recognized the Myc epitope located at after signal peptides (1-18 amino acids) of the WT and mutant receptors (**Figure S4 a,b**)

The full-length Ocn 1-49 peptide dose-dependently stimulated cAMP accumulation in the wildtype receptor, but the response was significantly decreased in the K352E, H355P and K352E/H355P mutant receptors (Figure 5A). Cells transfected with these three mutants hGPRC6A cDNAs also showed no response to Ocn 20-29 stimulation (**Figure 6A**). These results indicate that the binding sites identified by the computational modeling are accurate and that the helical region (352–363) is important for Ocn binding.

**Figure 6.**
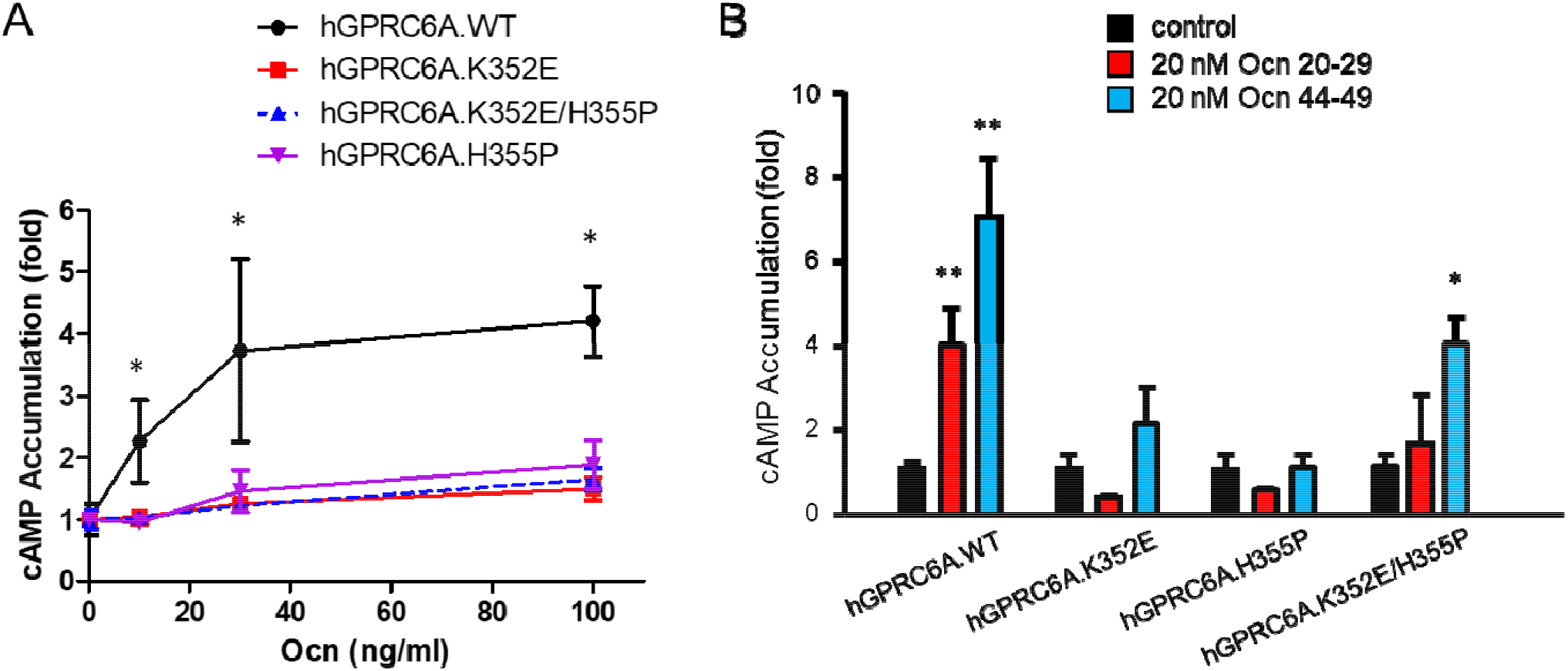
Mutagenesis of residues in predicted Ocn-binding pocket of GPRC6A. A, Comparison of dose-dependent effects of Ocn on cAMP accumulation in HEK-293 cells transfected with the cDNA plasmids of WT, K352E, H355P, or K352E/H355P hGPRC6A. * indicate a significant difference from control and stimulation groups at p<0.05 and p<0.01, respectively. B, Comparison of Ocn small fragments, Ocn 20-29 or Ocn 44-49 at 20 nM concentration on cAMP accumulation in HEK-293 cells transfected with the cDNA plasmids of WT, K352E, H355P, or K352E/H355P hGPRC6A. * and ** indicate a significant difference from control and stimulation groups at p<0.05 and p<0.01, respectively.

The Ocn 44-49 C-terminal fragment, which was predicted to bind to the 7TM in our prior homologous model (11), also had a significantly attenuated signaling response in the single mutants. Interestingly, Ocn 44-49 was able to partially activate the double mutant as assessed by cAMP accumulation (**Figure 6B**).

## Discussion

Recent Cryo-EM analysis of Family C mGluR5 (16) and CaSR (17,19,20) receptors describes the structural basis for their orthosteric ligand activation by calcium and L-amino acids and allosteric modulating effects of peptides and small molecules. In the current study, we developed structural homology models of GPRC6A based on the Cryo-EM and found evidence for both orthosteric and allosteric binding sites in its VFT, similar to mGluR5 and CaSR.

GPRC6A is activated by L-amino acids, cations, and the Ocn peptide (4). Prior work examined the possible binding of the hexapeptide C-terminal domain of Ocn and identified a binding site in the 7TM domain of GPRC6A for this fragment 44-48 (11). However, a comparison of recently available Cryo-EM structures of mGluR5 and CaSR suggested that a regulatory binding site in the VFT domains of Family C receptors may exist. These results led us to further examine the Ocn induced activation of GPRC6A and its isoform-specific modulation. Interestingly, GPRC6A isoforms only differ in the extracellular domain (i.e. VFT) which suggests that a second binding site for Ocn may exist in the VFT. Taken together, the new structural models developed for GPRC6A and complimentary experiments and peptide-protein docking calculations reported here provide clear evidence of a binding site for full-length and/or residues 20-29 Ocn within the GPRC6A VFT, located near the helical region (352–363).

Reports of GPRC6A activation by the peptide Ocn are inconsistent, with some studies showing activation (2,3,5,11) and other studies finding no effect (29). Here, we confirmed Ocn activation of GPRC6A in *in vitro* culture models. Moreover, the lowering of glucose and stimulation of IL-6 24 hours after a single injection of Ocn 20-29 in wild-type mice but not *Gprc6a*^-/-^ mice validate *in vivo* effects of this peptide mediated through GPRC6A. The failure to observe expected stimulation of insulin and adiponectin may be due to examining one time point, the short observation period or the limited doses tested. Regardless, the location of the Ocn binding site in the VFT at a site distinct from the orthosteric binding site suggests Ocn functions as a PAM rather than an orthosteric ligand. The fact that Ocn activation of GPRC6A requires the presence of calcium (3,5,11) *in vitro* is consistent with Ocn function as a PAM for GPRC6A (2,11).

Our findings also suggest altered ligand specificity and downstream signaling of GPRC6A isoforms created by alternative splicing. The loss of a segment of the VFT in isoform 2 resulted in the loss of responses to L-Arg as well as Ocn. In contrast, isoform 3 retains responsiveness to orthosteric activation but exhibits biased signaling to Ocn, as evidence by loss of ERK but preservation of cAMP signaling. Loss of this segment may alter the conformational changes leading to opening of binding sites in the intracellular TM 6 domain that is important for G-protein coupling. If so, different tissue distributions or the ratio of these isoforms would be expected to change orthosteric ligand and PAM sensitivities as well as alter coupling to downstream signaling pathways resulting in tissue-specific regulation mechanisms.

Our modeling data shows that most contacts between Ocn and GPRC6A are driven by the structural part of the Ocn peptide (3 helical structure). Peptide fragments of Ocn consisting of 1-7, 8-43, and 44-49 are released from bone (30). We tested the GPRC6A activation potential of multiple Ocn fragments to further identify the key regions of Ocn that interface with GPRC6A. The hexapeptide 20-29 most strongly activated GPRC6A, consistent with its location in the crucial region of the full-length Ocn used in our docking studies to the VFT helix region. Ocn 8-19 (apvpypdplepr) also activated GPRC6A (**Figure 3**). This upstream peptide is similar to the pentadecapeptide peptide wlgapvpypdplepr that is reported to activate GPRC6A *in vitro* and to improve fatty liver disease and insulin resistance after oral or intraperitoneal administration to a mouse model (31). When examining the activities associated with 44-49, it is important to recall that previous docking and mutagenesis studies suggested that the C-terminus of Ocn binds to the extracellular transmembrane domain of GPRC6A (11). This might explain the effects of the rfygpv, C-terminal peptide of Ocn to activate GPRC6A in setting of mutations to disrupt the Ocn binding domain in the VFT helix. Regardless, short peptides derived from Ocn might be developed for therapeutics to activate GPRC6A, similar to the 7 amino acid etacalcitide PAM for CaSR.

Another interesting observation is the greater potency of the peptide 20-29 compared to Ocn 1-49. Hypothetically this may be due to a potential disulfide bond present in the small peptide that creates a stable complementary conformation within the predicted binding site. Alternatively, competitive self-interactions between the domain and the disordered N-terminus of Ocn may inhibit long-lived binding conformations between Ocn and GPRC6A. Instead, flexibility of the N-terminal domain that result in N-terminal-VFT contacts may inhibit conformational changes along the dimer interface of the VFT domains.

Although we did not examine binding of other PAMs to GPRC6A in this study,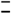testosterone and triphenol compounds that activate GPRC6A are predicted to bind to the 7TM domain of GPRC6A (12,13), analogous to the second PAM domain in the 7TM domain of mGluR5 and CaSR (16,20). This suggests that members of the Family C GPCRs, at least those with large VFT domains, have a conserved structure consisting of two allosteric binding sites, one in the VFT and the other in the 7TM domain, along with an orthosteric ligand binding domain in the VFT.

Finally, GPR158 is another proposed Ocn sensing receptor. The Cryo-EM structure of GPR158 has recently been defined (32). Our VFT Ocn binding region between GPRC6A and GPR158 is not conserved and a structure superimposition of the two structures shows that the helix region (352–363) identified for GPRC6A is missing in GPR158.

Our findings are limited by the lack of Cryo-EM structural data for GPRC6A. Cryo-EM analysis of mGluR5 and CaSR show that each functions as homodimers, where activation leads to closure of the VFT and conformational changes the 7TMs revealing G-protein binding sites on the cytoplasmic side necessary for signaling(17). For CaSR, which is most closely related to GPRC6A, receptor activation results in 7TM asymmetry in the homodimer leading to only one protomer for G-protein coupling being stabilized by PAM binding. Our biased signaling response to Ocn in isoforms 2 and 3 suggests that there may be a structural basis for the divergent cAMP and ERK responses to Ocn. To fully understand GPRC6A signaling we may need cryo-electron microscopy structures of GPRC6A in inactive or active states bound to the orthosteric ligands, PAMS and NAMs.

## Materials and methods

### System and docking details

GPRC6A and Ocn structures were modeled using AlphaFold2 (22,25,33), which is an AI system developed by DeepMind to predict a 3D structure of a protein from its amino acid sequence. Multiple sequence alignment (MSA) was performed using the Jackhammer method. The ‘max_recycles’ parameter, r which controls the maximum number of times the structure is fed back into the neural network for refinement, was set to 3. All the other parameters were set to default. The final models were ranked using the pLDDT score. Glycans were not added as the details are not well known. For docking, HADDOCK 2.4 (28) was used to dock the GPRC6A-VFP to Ocn. GPRC6A residue numbers 334,335,336,337 and mature Ocn residue numbers 21,22,23,24,25,26, and 27 were used to guide the docking. The best 4 models from the top cluster (containing 133 models) with HADDOCK score of −81.2±1.1 and Z-score of −1.5 were analyzed.

### Simulation details

Molecular dynamics simulation (MD) of the top ranked AlphaFold2 model was performed using the Amber simulation engine(34) to refine the structures. The MD was performed on the protein using Amber14 ff14SB (35) force field, with a non-bonded cutoff of 10 Å using the Particle Mesh Ewald algorithm (36). The protein-glycan bound system was hydrated by water model TIP3P (37) in an octahedral box of 10 Å around the protein in each direction. Initially, the protein was held fixed with a force constant of 500 kcal mol^-1^ Å^-2^ while the system was energy minimized with 500 steps of steepest descent followed by 500 steps with the conjugate gradient method. In a second minimization step, the restraints on the protein were removed and 1000 steps of steepest descent minimization was performed, followed by 1500 steps of conjugate gradient. The system was heated to 300 K while holding the protein fixed with a force constant of 10 kcal mol^-1^ Å^-2^ for 1000 steps. Then, the restraints were removed, and 1000 MD steps were performed. The SHAKE algorithm(38) was used to constrain all bonds involving hydrogen in the simulations. MD production runs were performed at 300 K using the NPT ensemble and a 2 fs time step. The temperature was fixed with the Langevin dynamics thermostat (39) and the pres sure was fixed with the Monte Carlo barostat (40). The last snapshot from the 20ns trajectory was used for all analyses.

### Measurement of Total and Phospho-ERK by ERK Elisa Analysis

Ocn was purified from bovine tibial bone extracts (41,42). Decarboxylated Ocn was produced by treating Ocn in vacuo at 110°C (42–44). The purity and decarboxylation state were confirmed by native gel electrophoresis (41), or by blotting followed by reaction with 4-diazobenzene sulfonic acid staining for γ-carboxyglutamic acid (42,43). The human Ocn fragments were synthesized by BioMatik USA (Wilmington, Delaware, USA).

All culture reagents were from Invitrogen (Waltham, MA, USA). Human embryonic kidney HEK-293 cells were obtained from American Type Culture Collection and cultured in DMEM medium supplemented with 10% fetal bovine serum and 1% Penicillin/Streptomycin (P/S). Briefly, HEK-293 cells transfected with/without human GPRC6A isoforms cDNA plasmids were starved by overnight incubation in serum-free DMEM/F12 containing 0.1% bovine serum albumin (BSA) and stimulated with various ligands at different doses. ERK activation was assessed 20□min after treatment by using ERK1/2 (phospho-T203/Y204) ELISA Kit (Invitrogen) corrected for the amount of total ERK using ERK1/2 (Total) ELISA Kit (Invitrogen) to measure ERK levels.

### Measurement of cAMP accumulation

HEK-293 transfected with human GPRC6A isoform cDNA plasmids cells (10^5^ cells/well) (45) were cultured in triplicate in 24-well plates in DMEM supplemented with 10% fetal bovine serum and 1% penicillin/streptomycin (100 U/mL of penicillin and 100 μg/mL of streptomycin) for 48 hours followed by 4 hours incubation in DMEM/F12 containing 0.1% BSA and 0.5 mM IBMX to achieve quiescence. Quiescent cells were treated with vehicle control, various ligands at concentration as indicated for 40 minutes at 37°C. Then, the reaction was stopped, and the cells lysed with 0.5 mL 0.1N HCl. cAMP levels were measured by using Cyclic AMP EIA kit (Cayman Chemical) following the manufacture’s protocol.

### Mouse and serum biochemistry

8 week old wild-type (*Gprc6a*^+/+^) and global knock out (*Gprc6a*^-/-^) mice were injected intraperitoneal with Ocn 20-29 (2 μM/kg body weight), or vehicle (saline; 10 μl/g body weight). Serum was collected at 24 hours after intraperitoneal injection. Blood glucose levels were measured by using blood glucose strips and the Accu-Check glucometer as described (7,27). Insulin (mouse) ultrasensitive ELISA kit was obtained from ALPCO Immunoassays (Salem, NH, USA). Mouse IL-6 ELISA Kit was purchased from Invitrogen (Waltham, MA, USA). Mouse Adiponectin PicoKine™ Quick ELISA Kit was obtained from mybiosource.com.

The *Gprc6a*^-/-^ mouse model was created by replacing exon 2 of the GPRC6A gene with the hygromycin resistance gene, as described previously (46). Mice were maintained and used in accordance with recommendations as described (National Research Council 1985; Guide for the Care and Use of Laboratory Animals Department of Health and Human Services Publication NIH 86-23, Institute on Laboratory Animal Resources, Rockville, MD) and following guidelines established by the University of Tennessee Health Science Center Institutional Animal Care and Use Committee. The animal study protocol was approved by the institutional review boards at University of Tennessee Health Science Center Institutional Animal Care and Use Committee.

### Site-directed mutagenesis

*In-vitro* mutagenesis by PCR-mediated recombination QuikChange II XL Site-Directed Mutagenesis Kit (Agilent) was performed using the cDNA of human GPRC6A cloned in the vector pcDNA3 (Invitrogen) as template. The primer sets as following. K352E.sen: GTGACAGTCACGAACTCTTACATG; K352E.antisen: CATGTAAGAGTTCGAGACTGTCAC; H355P.sen: CAAACTCTTACCTGAATATGCCATG; H355P.antisen: CATGGCATATTCAGGTAAGAGTTTG; K352E/H355P.sen: GTGACAGTCACGAACTCTTACGTGAATATGC; and K352E/H355P.antisen: GCATATTCAGGTAAGAGTTCGTGACTGTCAC. All mutants were confirmed by sequencing.

### Statistics

We evaluated differences between groups with the Student’s *t* test, and for multiple groups by two-way ANOVA, followed by *a post-hoc* Tukey’s test. Significance was set at p□<□0.05. All values are expressed as means ± SEM. All computations were performed using the Statgraphic statistical graphics system (STSC Inc., Rockville, MD, USA).

## Abbreviations

cAMP: cyclic adenosine monophosphate
CRD: cysteine rich domain
ERK: extracellular-signal-regulated kinase
GPRC6A: G protein-coupled receptor family C group 6 member A
NAM: negative allosteric modulator
Ocn: osteocalcin
PAM: positive allosteric modulator
VFT: venus flytrap
TM: transmembrane

## Acknowledgements

This work was supported by grants from the National Institutes of Health, NIAMS grant number 1R01AR071930 and NIDDK grant numbers 1R01DK121132 and 1R01DK120567 (LDQ) The content is solely the responsibility of the authors and does not necessarily represent the official views of the National Institutes of Health.

## Author Contributions

Rupesh Agarwal, Micholas Dean Smith and Jeremy C. Smith were responsible for the computational studies and Min Pi, Ruisong Ye, and L. Darryl Quarles were responsible for the experimental studies regarding GPRC6A. All authors contributed to the writing of the paper.

## Supporting Information

**Table S1.**
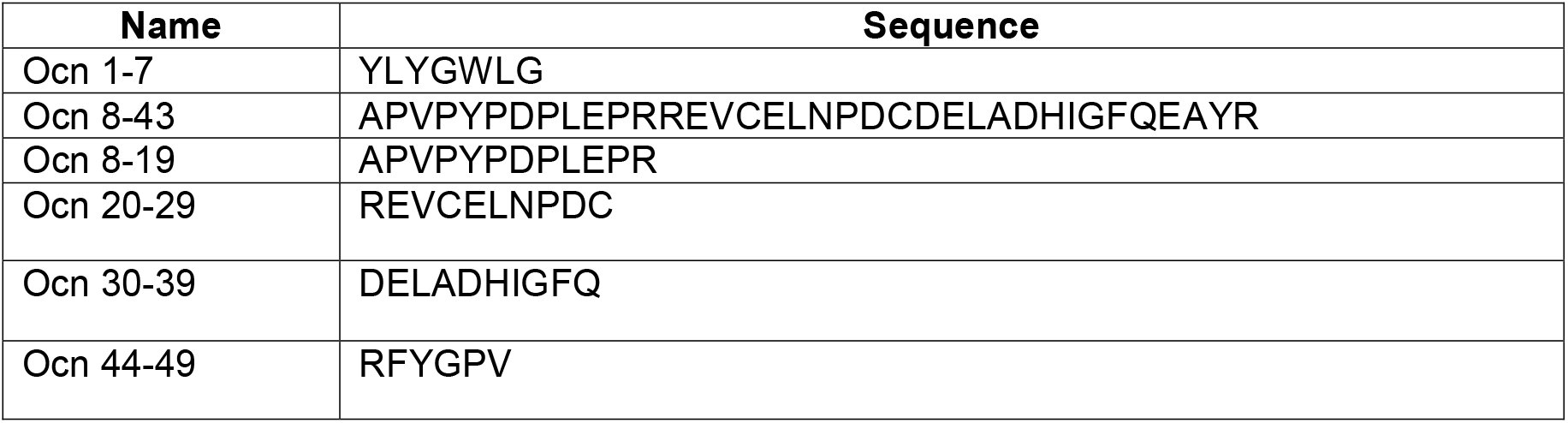
The amino acid sequence of synthesized human Ocn.

**Figure S1:**
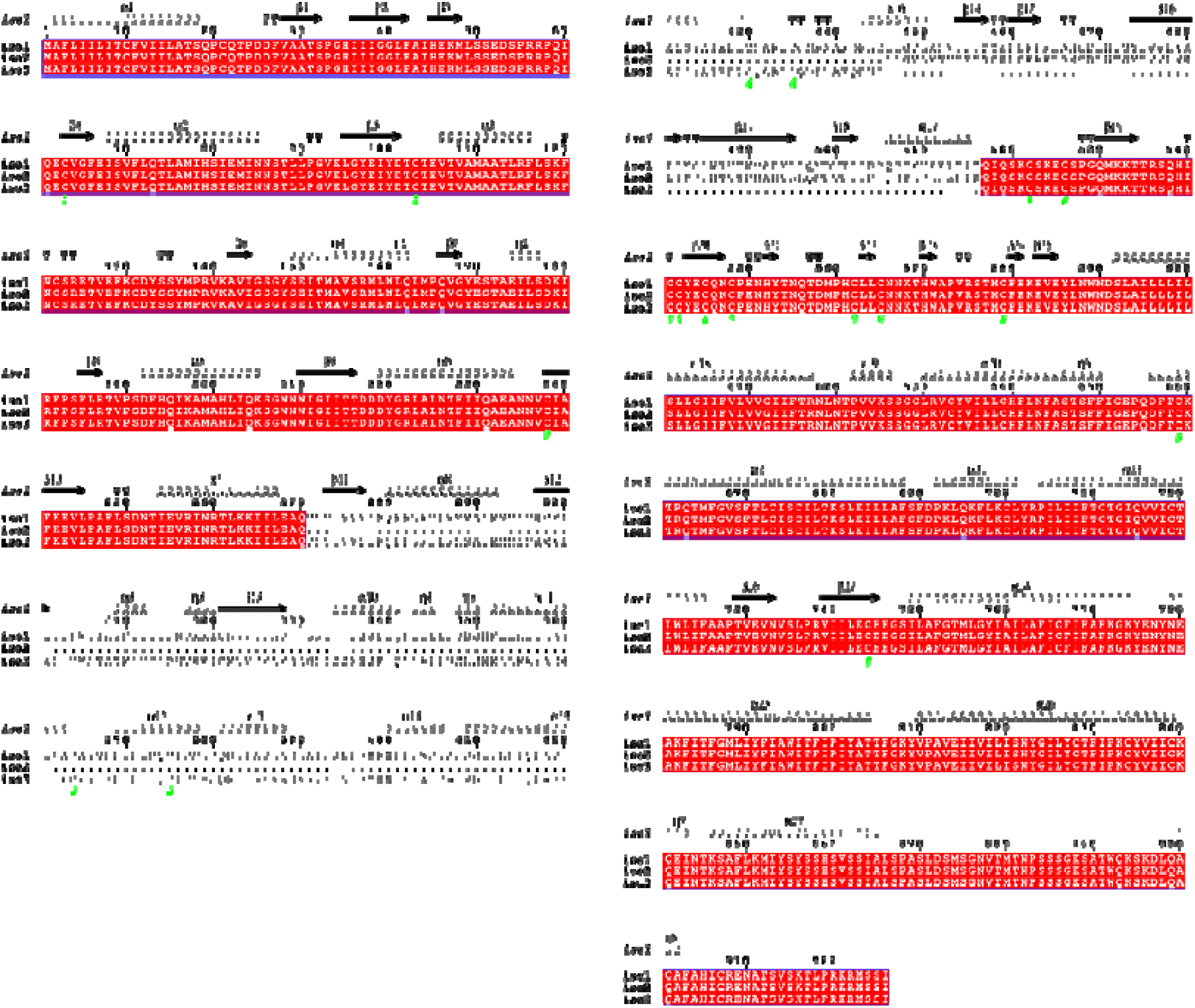
Sequence alignment of GPRC6A isoform 1, 2 and 3.

**Figure S2:**
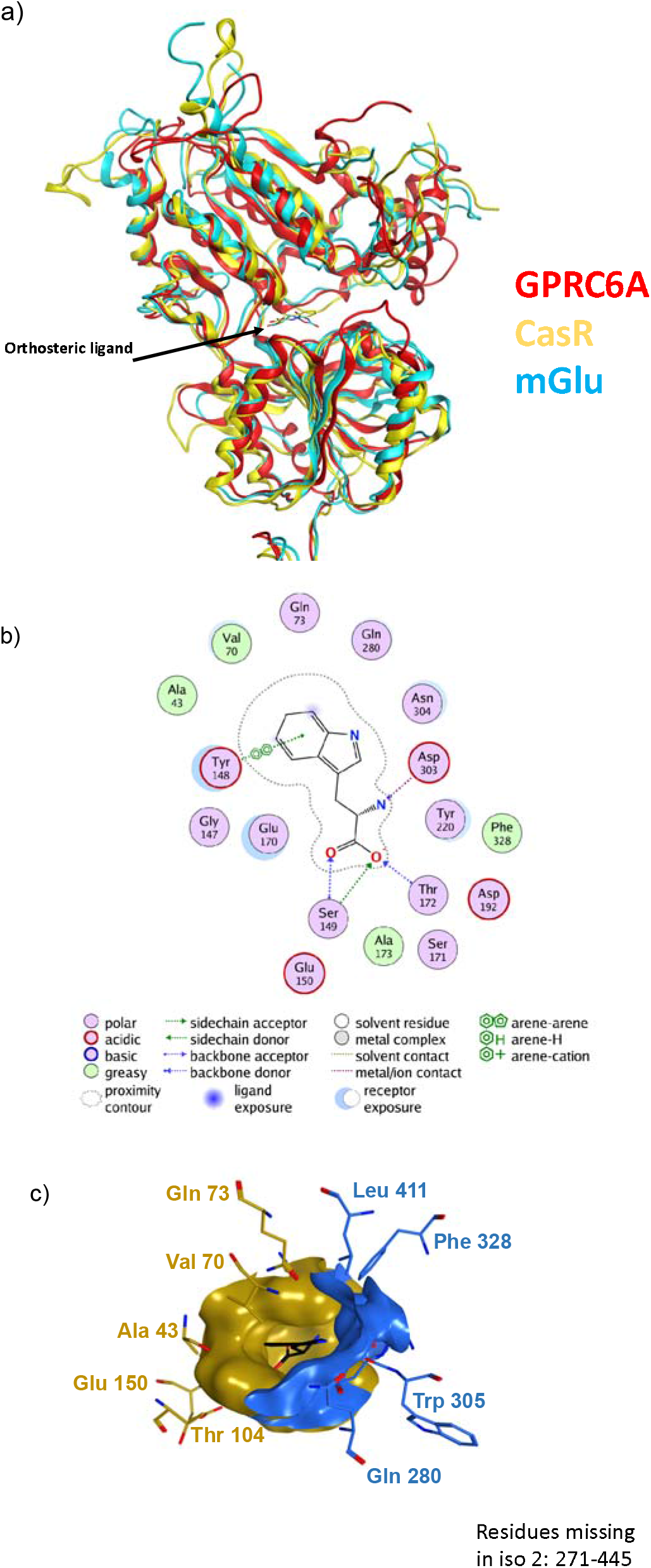
a) Superimposed structure of AlphaFold2 minimized model of GPRC6A, CaSR with L-Trp (PDB 7DTU) and mGluR5 with orthosteric analogue L-quisqualate (PDB 6N51) (in ribbon) showing orthosteric ligands. b) Interaction map of L-Trp in isoform 1 c) Modeling of the orthosteric ligand from CaSR in GPRC6A-isoform 1. Black: Trp (placed using CaSR:Trp as template); Blue (orthosteric residues/ orthosteric binding pocket region missing in iso2); yellow: iso1 orthosteric pocket region.

**Figure S3:**
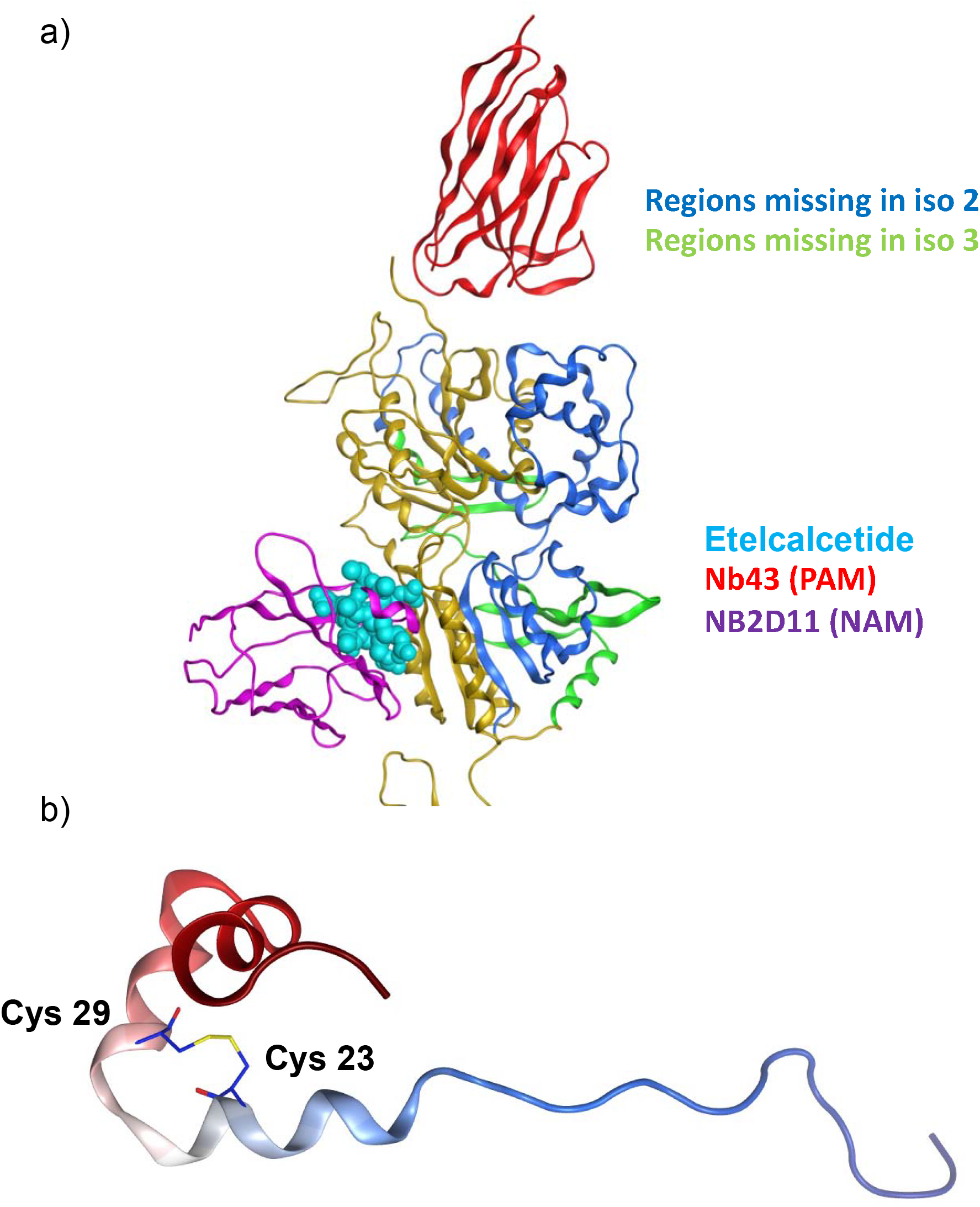
a) Superimposition of crystal structures on GPRC6A model. Yellow ribbon: GPRC6A; Dark blue ribbon: region missing in iso2; green ribbon: part missing in iso3; Red ribbon: PAM Nb43 from mGluR5 (PDB 6N4Y); magenta ribbon: NAM NB2D11 from CaSR (PDB 7E6U); Cyan sphere: etelcalcetide (calcimimetic drug) (PDB 7M3G); b) AlphaFold2 minimized model of hOcn colored from N to C terminus (Blue to red) showing disulfide bond between Cys23 and Cys29.

**Figure S4.**
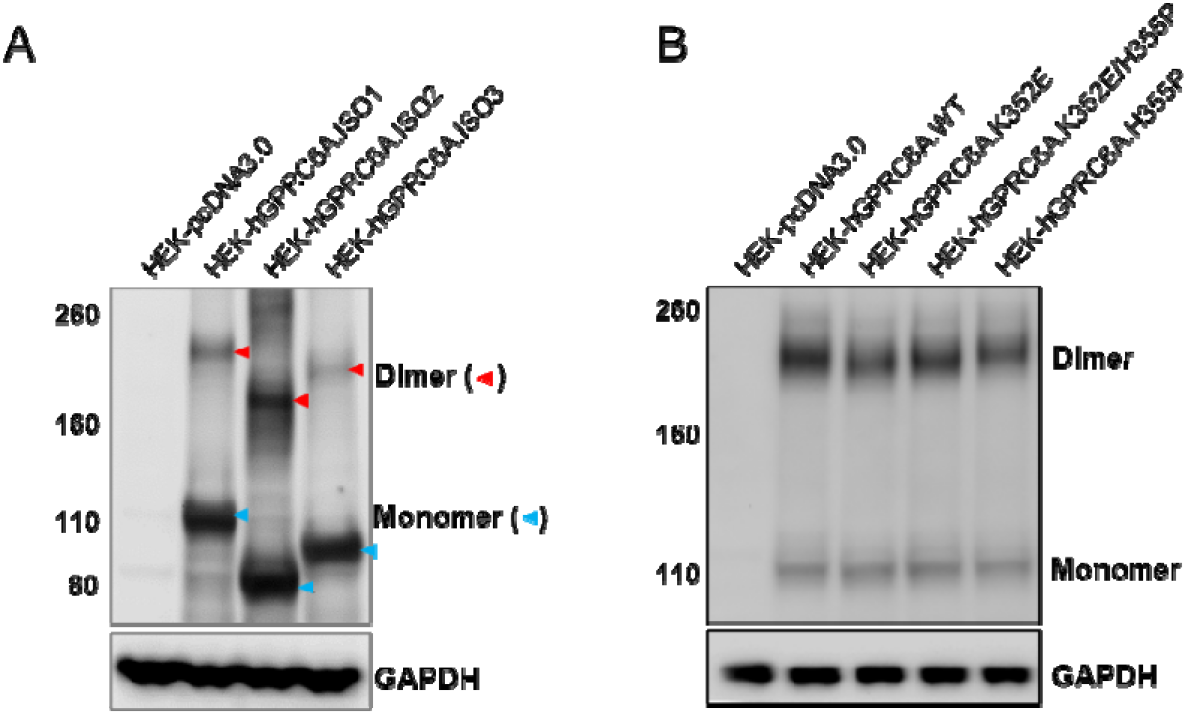
The protein expression of GPRC6A isoforms (A) and mutants (B).

**Figure S5.**
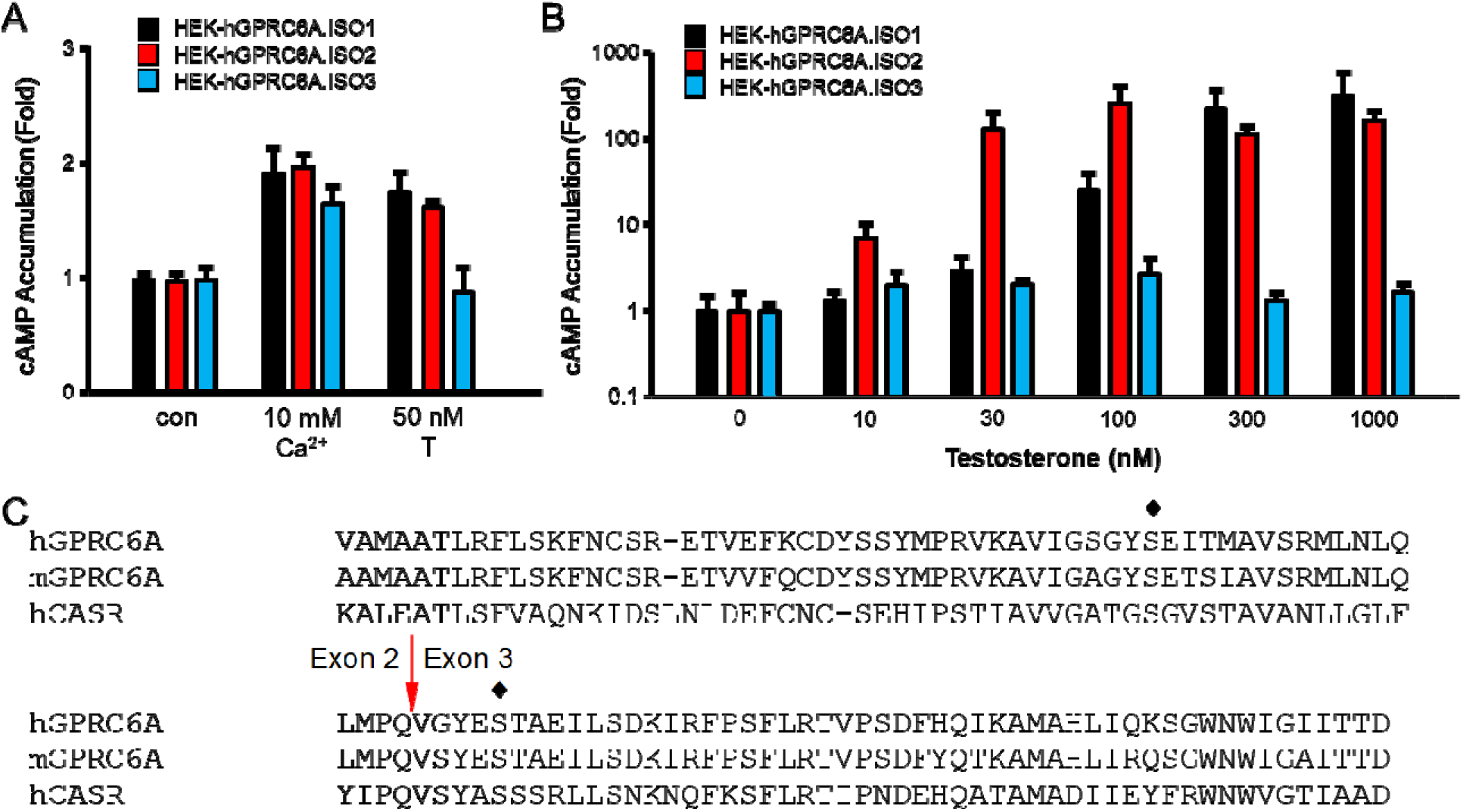
GPRC6A isoform 3 loss responsive to T activation. A. Comparison of effects of Ca^2+^ or T on GPRC6A isoforms mediated cAMP accumulation. B. Comparison of dose-dependent effects of T on GPRC6A isoforms mediated cAMP accumulation. HEK-293 cells were transfected with cDNA plasmids of GPRC6A isoform 1, 2 or 3 for 48 hours, after incubated in Dulbecco’s modified Eagle’s medium /F-12 containing 0.1% bovine serum albumin quiescence media for 4 hours, then exposed to Ca^2+^ or T at indicated concentrations for 40 minutes for cAMP accumulation details as described under “Methods”. * and ** indicate a significant difference from control and stimulation groups at p<0.05, and 0.01. C. Alignment of human GPRC6A, mouse GPRC6A and human CaSR showing predicted Ca^2+^ binding sites. Red arrow shows the junction of exon 2 and exon 3. ♦ indicate expected amino acid in GPRC6A depended by CaSR.

